# In situ Patch-seq analysis of microglia reveals a lack of stress genes as found in FACS-isolated microglia

**DOI:** 10.1101/2023.03.22.533782

**Authors:** Olga Bakina, Thomas Conrad, Nina Mitic, Ramon Oliveira Vidal, Tessa Obrusnik, Somesh Sai, Christiane Nolte, Marcus Semtner, Helmut Kettenmann

**Affiliations:** Cellular Neurosciences, Max-Delbrück Center for Molecular Medicine, Berlin, Germany; Humboldt Universität, Berlin, Germany; Genomics platform, BIMSB, Max-Delbrück Center for Molecular Medicine, Berlin, Germany; Quantitative Developmental Biology, Berlin Institute for Medical Systems Biology, Max Delbrück Center for Molecular Medicine, Berlin, Germany; Institute of Chemistry and Biochemistry, Department of Biology, Chemistry and Pharmacy, Freie Universität Berlin; Shenzhen Institute of Advanced Technology, Chinese Academy of Sciences Shenzhen, China

**Author notes:** **For correspondence:** Prof. Dr. Helmut Kettenmann, Cellular Neurosciences, Max Delbrueck Center for Molecular Medicine in the Helmholtz Society Robert-Roessle-Strasse 10, 13125 Berlin, Germany. These authors contributed equally to the manuscript (Marcus Semtner, Helmut Kettenmann).

**Keywords:** microglia, mouse, brain, FACS, Patch-seq, electrophysiology, single-cell RNA-sequencing, tissue-dissociation stress response

## Abstract

We applied the patch-seq technique to harvest transcripts from individual microglial cells from cortex, hippocampus and corpus callosum of acute brain slices from adult mice. After recording membrane currents with the patch-clamp technique, the cytoplasm was collected via the pipette and underwent adapted SMART-seq2 preparation with subsequent sequencing. On average, 4138 genes were detected in 113 cells from hippocampus, corpus callosum and cortex, including microglia markers such as *Tmem119, P2ry12* and *Siglec-H*. Comparing our dataset to previously published single cell mRNA sequencing data from FACS-isolated microglia indicated that two clusters of cells were absent in our patch-seq dataset. Pathway analysis of marker genes in FACS-specific clusters revealed association with microglial activation and stress response. This indicates that under normal conditions microglia *in situ* lack transcripts associated with a stress-response, and that the microglia-isolation procedure by mechanical dissociation and FACS triggers the expression of genes related to activation and stress.

## Introduction

Microglia are the resident macrophages of the central nervous system. They originate from myeloid progenitors in the embryonic yolk sack and invade the developing brain before the formation of the blood-brain barrier (Perdiguero et al. 2015). They migrate into all regions of the central nervous system including spinal cord, retina and optic nerve. They are also present in both white and grey matter. In the adult normal brain microglia transform from an ameboid morphology as observed in the immature brain into a highly ramified phenotype, termed ‘resting’ microglia in the classic literature. In the last years it has been established, however, that these ramified cells are constantly surveying the environment by rapid movements of their processes in the normal brain (Davalos et al, 2005, Nimmerjahn et al 2005). During any type of pathologic event, microglia transform into an activated state. This activated state can be highly diverse and microglia can express and release cytokines and chemokines, neurotrophic and growth factors by which they influence the pathologic process (Hanisch and Kettenmann, 2007). A series of recent studies has applied single-cell RNA sequencing (scRNA-seq) to identify microglial phenotypes in different brain regions and different developmental and pathological states (Masuda et al. 2019; Hammond et al. 2019; Keren-Shaul et al. 2017). Distinct populations of microglia were identifed both in the developing normal brain and under pathological conditions. For these studies, microglia were isolated from brain tissue using mechanical or enzymatic dissociation and subsequently purified either by FACS or by magnetic columns.

In the present study, we applied a technique which avoids cell isolation and sorting, the *in situ* patch-seq procedure, to characterize individual microglia in acutely isolated brain slices. Thus, the cell remains in its tissue context while the RNA is harvested. This technique was first developed to correlate the expression profile of neocortical neurons with their electrophysiological profile (Cadwell et al., 2016; Fuzik et al., 2016). A whole cell recording configuration is established with the patch-clamp technique which allows access to the cytoplasm of the cell via a glass pipette. By applying negative pressure to the pipette, the cytoplasm of that single, individual cell can be harvested and the RNA can be sequenced (Cadwell et al., 2017). Here, we applied this technique to microglia in three distinct regions of the brain, hippocampus, cortex and the corpus callosum and compared our single cell profiles to those obtained after cell isolation and FACS sorting. Our results indicate that the stress response previously observed in isolated microglia from wild type mice may have been triggered during tissue disruption and cell sorting, and does not reflect the normal state *in situ*.

## Results

### A PATCH-SEQ PROTOCOL FOR HARVESTING RNA FROM SINGLE MICROGLIAL CELLS IN BRAIN SLICES

To collect RNA from single microglial cells via the patch clamp pipette, we used 250 μm thick brain slices of 10-14 weeks old transgenic mice expressing EGFP under control of the colony stimulating factor 1 receptor promoter (“Mac Green” mice (Sasmono et al. 2003) (**Fig. 1A**)). Microglial cells were identified by their EGFP fluorescence and they displayed the typical ramified morphological features of microglia as previously described (Boucsein, Kettenmann, and Nolte 2000). Microglia from different brain areas, namely cortex (CX), hippocampus (HC) and corpus callosum (CC) were approached by a patch pipette (**Fig. 1B**). After establishing the whole cell recording mode, a series of de- and hyperpolarizing voltage steps ranging from -170 to +60 mV from a holding potential of -70 mV was applied. Additionally, we collected negative controls (<u>e</u>xtracellular <u>c</u>ontrols, EC) to control for the non-specific contamination of microglia samples with extracellular material by applying the same steps as for the sample collection but omitting to approach any cell (see Methods for details). We refer to all the collected samples including EC as “samples”, while “cells” are only samples from microglial cells.

**Figure 1.**
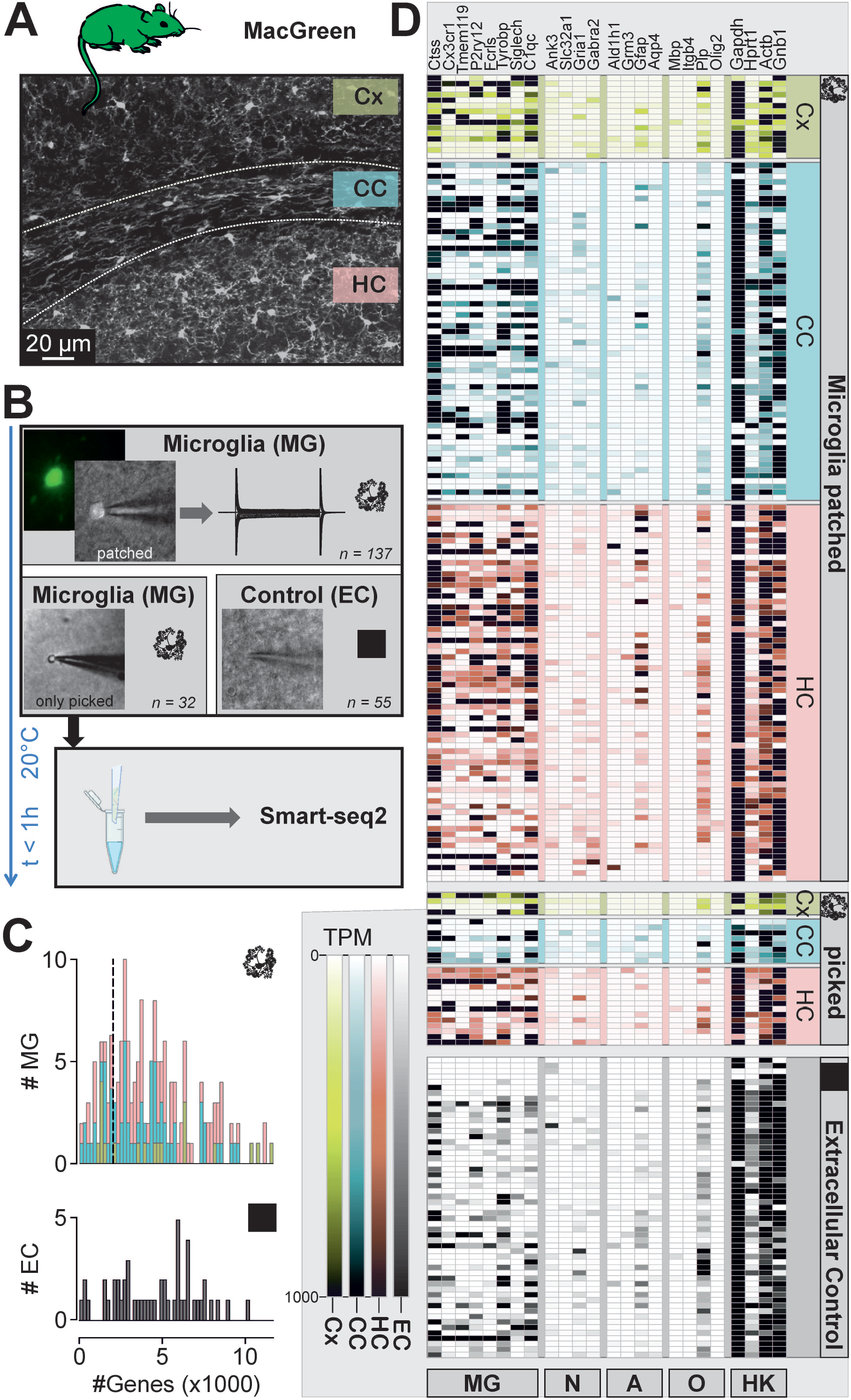
Patch-seq of microglia: method development and data quality analysis. **A** Confocal Image of a brain slice generated from a MacGreen mouse as used in this study, showing IBA^+^ microglial cells (immunolabeled as described in Matyash et al 2017)). Investigated regions-cortex (Cx), hippocampus (HC) and corpus callosum (CC) are marked. **B** Experimental scheme. (Upper image) Microglial cells (MG) were identified by their EGFP flurorescence (small fluorescence image on the left) and approached with the patch-clamp pipette to establish the whole cell configuration (phase contrast image, patched). A series of de- and hyperpolarizing voltage steps ranging from -170 and +60 mV were recorded from each cell as shown on the right. (Middle images) The cytoplasm was harvested by suction and subsequently the pipette was withdrawn. In some cases, the soma remained attached as shown in the left image. Extracellular controls (EC) were generated by moving a patch pipette 20 μm to 50 μm below the slice surface without approaching a cell. (Lower image) The content of the patch pipettes was expelled into a tube and Smart-seq protocol was immediately applied. Number of MG samples (137 patched, 32 only picked from cell-attached mode, 55 Extracellular controls) are indicated. **C** Histograms depicting the number of genes (x axis) obtained in MG (*top*) and EC (*bottom*) samples (y axis). **D** Heatmap displaying the expression of selected microglial (MG), neuronal (N), astrocytic (A), oligodendrocytic (O) marker genes as well as some housekeeping genes (HK). Brain regions are indicated on the right: cortex (Cx), hippocampus (HC) and corpus callosum (CC). The samples collected for RNA assessment were either microglial cells which were patched (Microglia patched), just picked while the electrophysiological profile was not recorded (picked), or extracellular controls.

We developed an optimized procedure to obtain *in situ* transcriptome information from the recorded cell. Subsequent to recording of membrane currents, a negative pressure of -70 mbar was applied for 5-10 min to the patch-pipette to harvest the cytoplasmic content of the cell. The pipette content was transferred to the lysis buffer followed by SMART-seq2 protocol for single cell cDNA library preparation (**Fig. 1B**). In approximately 50% of cases upon removing the pipette from the slice, the microglial soma was attached to the pipette and was ejected to the lysis buffer (**Fig. 1B**). In a portion of cells (n=28), the break-in was not performed after forming a Gigaohm seal, and the cells were collected from cell-attached mode with the pipette for RNA assessment. In order to ensure optimal cDNA quality, we performed a series of pilot experiments to optimize the protocol for cell harvesting and cDNA library preparation. Comparison of Tapestation profiles of samples which were snap-frozen in liquid nitrogen after collection with samples which were immediately processed further after collection showed that direct fresh processing resulted in a higher cDNA quality **(Suppl. Fig. 1)** and yielded ∼6x more detected genes (data not shown). Thus, all the samples collected for further data analysis were freshly prepared within maximally 1 hour from collection to the start of SMART-seq2 reaction while being kept at 4°C. In total, 244 samples were collected via patch-seq, of which 189 were cell samples collected from the three different regions, hippocampus (94), corpus callosum (76) and cortex (19) whereas 55 were EC. The average number of genes that were identified in these samples was 4289 ± 200 for cell samples and 4622 ± 320 for EC which was not significantly different (p > 0.9999) **(Fig. 1C)**, and suggesting that maneuvering the patch pipette below the slice surface produces off-target mRNA collection. To assess the general quality of our single cell transcriptomic dataset, we calculated TPM (transcripts per million) values and mapped the gene expression of classical markers for microglia (Ctss, Cx3cr1, Tmem119, P2ry12, Fcrls, Tyrobp, Siglech, C1qc), neurons (Ank3, Slc32a1, Gria1, Gabra2), astrocytes (Ald1h1, Grm3, Gfap, Aqp4) and oligodendrocytes (Mbp, Itgb4, Plp, Olig2) as well as housekeeping genes (Gapdh, Hprt1, Actb, Gnb1) from two recent surveys of mouse cortical diversity (Tasic et al., 2016; Zeisel et al., 2015). As evident from this heatmap **(Fig. 1D)**, one can appreciate that most of the microglial samples strongly express at least one of the microglia-specific genes, however, with some neuronal, astrocyte and oligodendrocyte markers being also present. Of note, in EC samples which should ideally feature no genetic material, there were apperently as many housekeeping genes as in cellular samples, suggesting that RNA sampling occurs to a certain extent just by moving the patch pipette through the tissue and thereby passing cellular entities. EC also contained microglial markers, however, at an apparently lower level than microglial samples.

### ELECTROPHYSIOLOGICAL PROPERTIES OF MICROGLIA IN CX, HC and CC

We compared the membrane currents of microglia from cortex, hippocampus and corpus callosum (**Fig. 2**). Consistent with previous data from us and many other labs (for reviews see Kettenmann *et al*. (2011)), microglial cells displayed characteristic membrane currents in all investigated regions when repetitively clamped at potentials between -170 and +60 mV starting from a holding potential of -70 mV (**Fig. 2A** and **B**). Cells were characterized by high input resistances and a small inwardly-rectifying potassium conductance in the range between -40 and -170 mV. Series resistances were similar throughout our recordings (**Fig. 2B**), providing a good comparability of microglia recordings from different brain areas. The current to voltage relationships of microglia from the observed brain regions were, however, not significantly different. Membrane capacities (**Fig. 2C**) were significantly higher in hippocampus (22.04 ± 0.79 pF, n = 40) as compared to corpus callosum (16.2 ± 0.91 pF, n = 38; p < 0.0001), and to cortex (17.6 ± 1.6 pF; n = 13; p = 0.0290) which might reflect morphological differences in these brain areas. There were no significant differences in membrane resistances (p = 0.0900; **Fig. 2C**). Reversal potentials were more depolarized in corpus callosum (−17.4 ± 1.5 mV) which was significantly different to hippocampus (−26.6 ± 1.9 mV; p = 0.0007) but not to cortex (−23.7 ± 2.8 mV; p = 0.1669).

**Figure 2.**
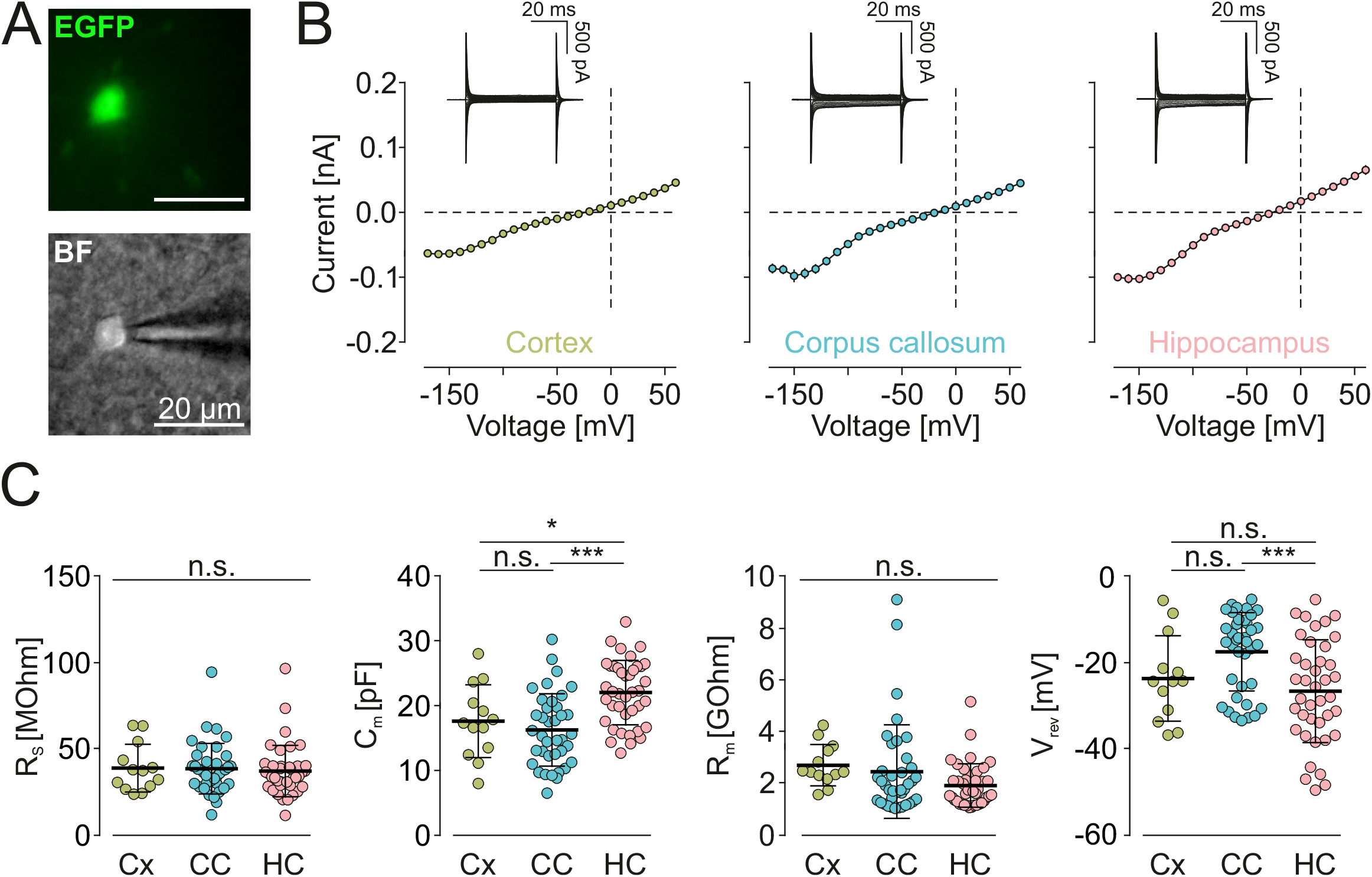
Electrophysiological data obtained before harvesting the cytoplasm for sequencing. **A** EGFP Fluorescence (top) and brightfield image (BF, bottom) of a microglial cell approached by a patch pipette. **B** (Inset) Samples of membrane currents from microglial cells clamped for 50 ms to a series of de- and hyperpolarizing voltages ranging from -170 mV to +60 mV from a holding potential of -70 mV. Cells were recorded in cortex, hippocampus and corpus callosum as indicated. (Main) Based on the recorded membrane currents, average current-voltage relationships for all patch-clamped cells in the notified region were constructed. **C** Summary of series resistances (R_s_; *left*), capacities (C_m_; *middle left*), membrane resistances (R_m_; *middle right*) and reversal potentials (V_rev_; right) of microglia from cortex (Cx), corpus callosum (CC) and hippocampus (HC). Significant differences are indicated by *** (p<0.001) and * (p=0.029), non-significance is indicated by n.s.

### QUALITY CONTROL OF PATCH-SEQ SAMPLES

Contamination of the samples with off-target cell transcripts is a characteristic feature of Patch-seq datasets from *in vivo* or *in situ* experiments (Tripathy, 2018). In order to explore the quality of collected transcriptomic samples, we applied UMAP clustering analysis to the dataset of cellular and EC controls. Clustering was based on top 80% variable genes across the dataset. Primary to further processing of the data, the following filtering for quality control was applied (**Fig. 3A**): We only used data from microglia samples with more than 1000 detected genes and from those that did express at least one of the following established microglia markers: Cx3cr1, Tmem119, P2ry12, Fcrls, Olfml3, Gpr34, Siglech, Gpr84, Trem2, Socs3, Ccl2, Tyrobp, Ctss and C1qc, resulting in the exclusion of 57 microglia samples from further analysis. As shown in **Fig. 3B** (and **Suppl.Fig.4**), UMAP clustering led to the separation of the majority of microglial samples (Cluster 2: 87 MG and 11 EC) from the EC samples (Cluster 1: 26 MG and 40 EC), indicating that the transcriptomes from cellular material were indeed different from extracellular controls. EC samples that are not clearly separated from other microglia samples in Cluster 1 most likely occurred due to method-specific contamination of the pipet with the surrounding material upon collection of the cytosols. This contamination phenomenon has been previously described in the study of the Pavlidis laboratory with the computation evaluation of contamination in multiple patch-seq experiments (Tripathy et al., 2018).

**Figure 3.**
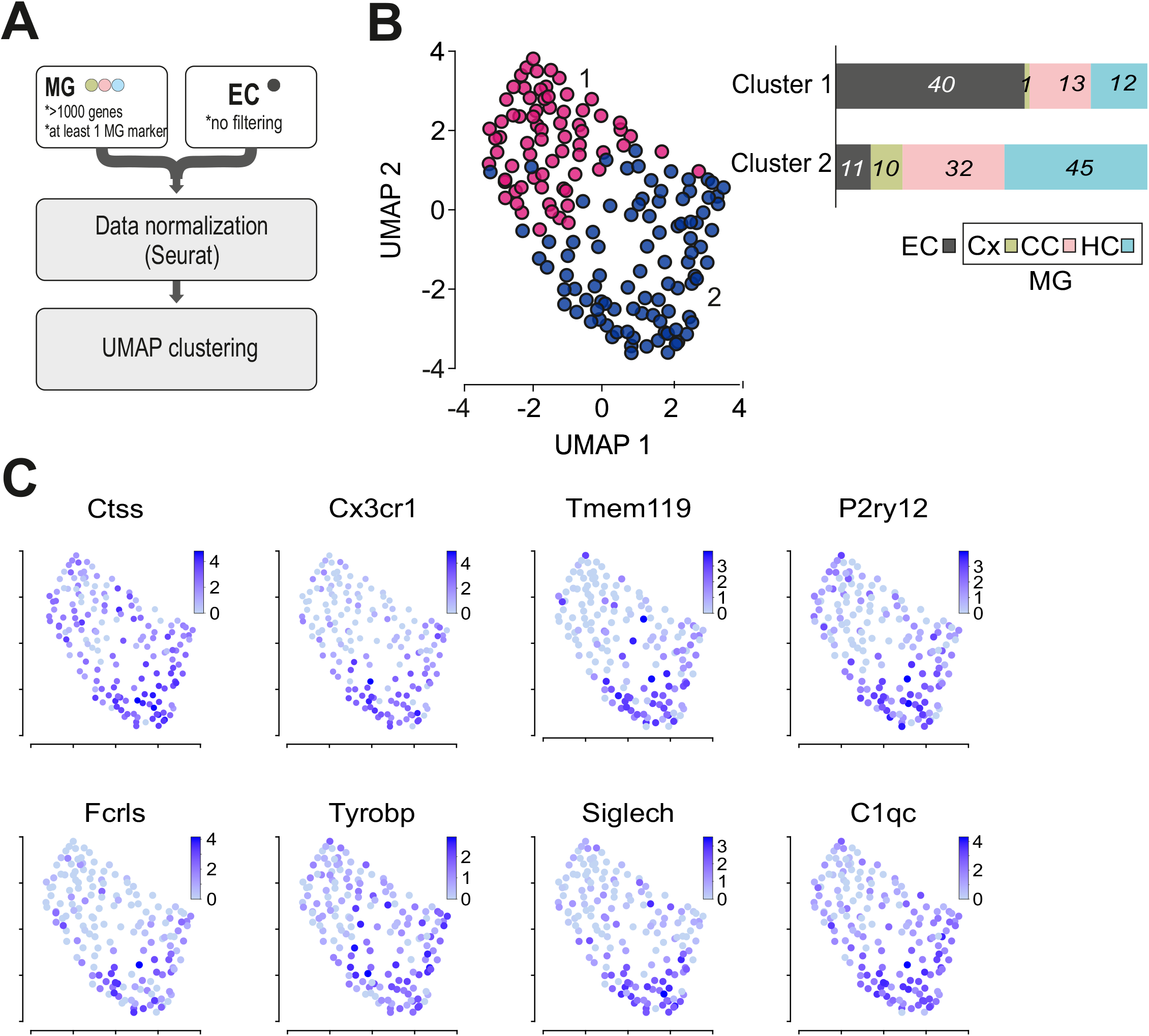
UMAP representation of MG and EC samples **A** Scheme depicting the filtering and computational pre-processing applied to the data set. **B** UMAP plot of MG and EC samples on top 80% variable genes. Clustering analysis revealed two clusters which are indicated by pink (Cluster 1) and purple (Cluster2) colors. As seen in the bar plot the majority of microglia samples are enriched in cluster 2, while the majority of EC samples in cluster 1. In the bar plot the y axis represents proportions of samples per cluster. Numbers on the plot indicate the number of samples in each cluster (EC (extracellular control, gray), MG (microglia) in Cx (cortex, green), CC (corpus callosum, pink), HC (hippocampus, blue).). **C** Expression levels of the microglia marker genes *Ctss, Cx3cr1, Tmem119, P2ry12, Fcrls, Tyrobp, Siglech* and *C1qc*. Note that all of these genes exert higher expression levels in Cluster 2.

While all of the cells expressed at least one of the known microglia markers, the vast majority of the cells expressed several of the microglia markers *Ctss, Cx3cr1, Tmem119, P2ry12, Fcrls, Tyrobp, Siglec-H, C1qc* (**Fig. 3C**). Based on UMAP analysis, there were no major differences found between hippocampus, corpus callosum and cortex samples, either supporting previous findings of a rather homogeneous microglial population in the adult mouse brain under normal conditions (Hammond et al., 2019; Masuda et al., 2019) or indicating that the sensitivity of the patch-seq method is too low for a proper brain region-dependent discrimination of microglial transcriptomes.

### A STRESS-ASSOCIATED SUBSET OF MICROGLIA AS REVEALED IN FACS SAMPLES IS NOT DETECTED IN THE PATCH SEQ DATA SET

As a next step we addressed the question of how our data set compares to previously published single cell RNA-Seq data sets of microglia isolated by mechanical dissociation of the tissue and FACS from corpus callosum and hippocampus (Masuda et al. 2019). In that study, microglia samples were isolated by dissociation of the brain in a glass potter, followed by 30 min Percoll centrifugation and a washing step. Subsequently, samples were stained with FACS antibodies, followed by cell sorting into 96 well plates with flow cytometry and SmartSeq2. We co-clustered our data (Patch-seq) and data from FACS-isolated microglia (Masuda et al., 2019) after log-normalization and integration using Seurat. Clustering analysis resulted in the identification of four clusters in the FACS dataset of which only two were also present in our Patch-seq datasets (Clusters 1 and 4), while two other clusters were specific for FACS-isolated cells (Clusters 2 and 3, **Fig. 4A**).

**Figure 4.**
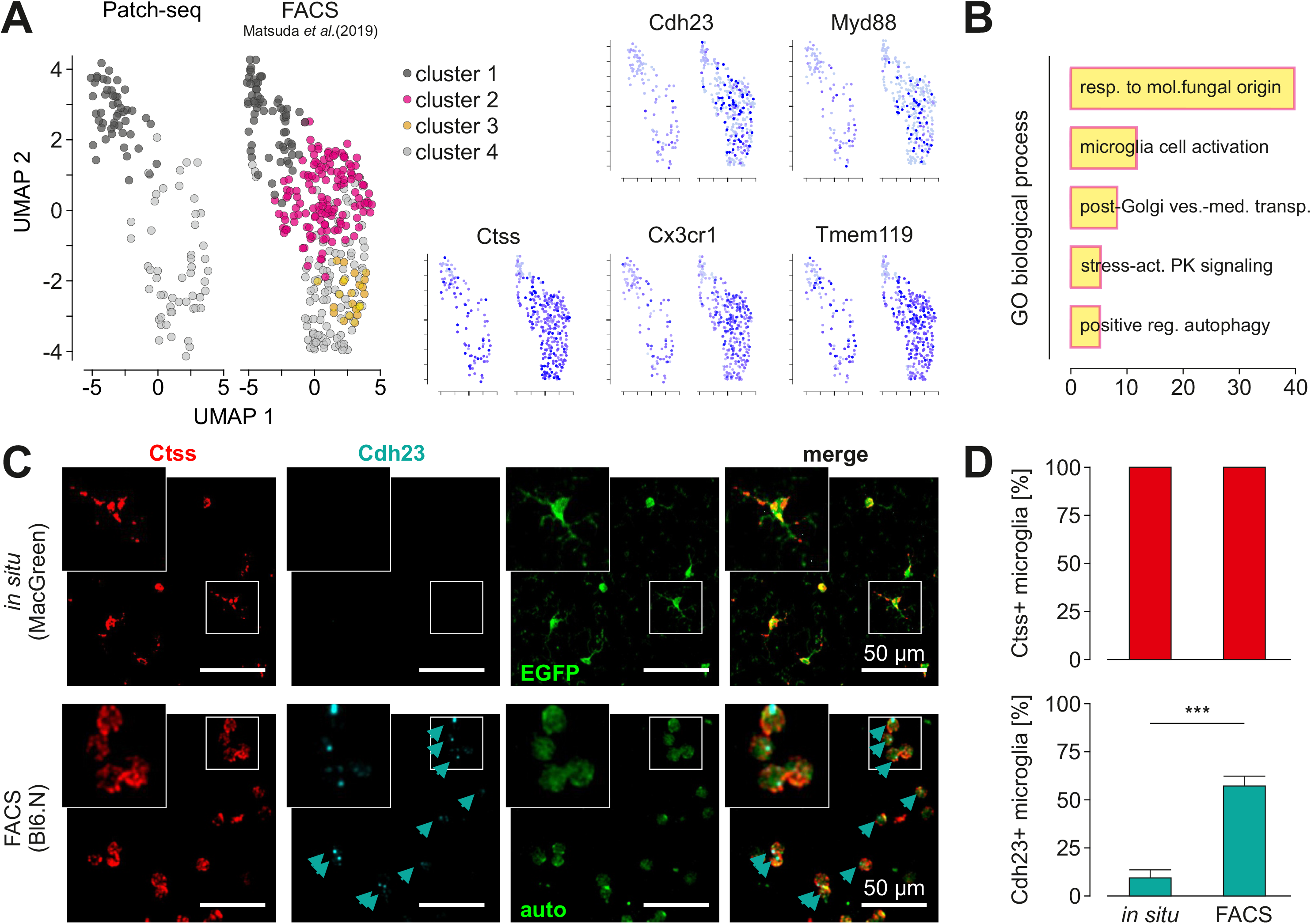
Comparison of single cell transcriptomes obtained by patch seq and after FACS sorting. **A** *Left*, Comparison of microglia single cell data from the current patch-seq and a previously published data set after FACS isolation (Matsuda et al., 2019). The latter data were obtained from microglia after brain dissociation and FACS. Note that clusters 1 and 4 appear in both data sets whereas clusters 2 and 3 are specific for FACS samples. *Right*, Gene expression analysis of the data sets shown in A for the microglia activation markers *Cdh23, Myd88* as well as for the homeostatic genes *Ctss, Cx3cr1* and *Tmem119*. **B** Gene ontology (GO) enrichment analysis of 275 top marker-genes of a FACS specific Cluster 2 reveals Cluster specific pathways. The bar chart represents the top 5 significantly enriched pathways in Cluster 2 in comparison to all other clusters. X-axis shows the fold enrichment of each pathway. **C** Detection of Ctss and Cdh23 in cortical brain slices from MacGreen mice (*top*) and FACS-isolated microglia from Bl6 mice (*bottom*) by RNAscope and confocal microscopy. Ctss (*left*) and Cdh23 (*middle left*) are indicated in red and cyan, respectively. Intrinsic EGFP signal of cell in slices is given in green (*middle right*); FACS isolated microglia in lower image show autofluorescence (auto). Merged images (*right*) show overlap of signals, indicated by arrows. The small square in each image is shown enlarged in the insets on the top left of each image. Scale bars: 50 μm. **D** Quantification of Ctss and Cdh23 in microglial cells in situ and after FACS isolation. Note that the homeostatic microglial marker Ctss (*left*) was expressed in both, *in situ* and *in vitro* microglial cells whereas Cdh23 (*middle*) was significantly more present after FACS sorting.

To characterize the gene signature of Cluster 2, we performed pathway analysis using the PANTHER gene overrepresentation test for the top 275 signature genes of Cluster 2 (p<0.01). Fisher’s exact test, with the Benjamini–Hochberg false discovery rate (FDR) correction for multiple testing was used for the over-representation test (Mi et al. 2019). This analysis revealed pathways involved in microglial cell activation: *C5ar1, Tlr2, Jun, Cx3cr1, Casp1*; stress-activated protein kinase signaling cascade: *Syk, Myd88, Ptpn1, Map3k3, Trib1, Taok2, Mapk8ip3, Errfi1*; and positive regulation of autophagy: *Flcn, Ulk2, Rab3gap1, Sptlc2, Tlr2, Fyco1, Atg16l1, Irgm2* (**Fig. 4B**), suggesting that Cluster 2 is associated with a stress response and microglia activation. Marker genes from Cluster 3 (another FASC-specific Cluster) are associated with autophagy (Tbc1d14), acidosis-induced cell death by chloride influx and cell swelling (Tmem206), metabolic repurposing of mitochondria (Sdha) (Mills et al., 2016), and therefore again with non-homeostatic functions. We tested the expression level of stress-related genes from Cluster 2 *in situ* in normal brain tissue and FACS-isolated microglia by single molecule fluorescent in situ hybridization (smFISH) for *Cdh23*. As positive control we used *Ctss*, a marker gene of clusters 1 and 4 (**Fig. 4C**). Cortical microglia from MacGreen mice, identified by their transgenic EGFP expression, were in 100% of cells positive for Ctss, whereas out of the 272 Ctss-positive cells in intact brain slices less than 5% expressed Cdh23 (**Fig 4C** and **D**). In contrast, 55% of microglial cells out of the total of 369 isolated from the brain using mechanical dissociation and FACS, expressed Chd23 (**Fig. 4C** and **D**), confirming that Cdh23, as representative gene from the transcriptomic signature of stress-associated Cluster 2, is indeed associated with the FACS-based single cell isolation procedure and is not normally expressed *in situ*.

## Discussion

In the present study, we compared the expression profile of microglia in the tissue context with microglia purified from the brain as single cells. For the former sample, we used the Patch-seq technique to harvest microglial RNA from cortex, corpus callosum and hippocampal tissue. This technique has been developed to analyze the expression profile of neuronal populations in the tissue context and compare it with the electrophysiological properties of these neurons (Cadwell et al., 2016, 2017). Here, the protocol has been adapted and applied to study microglial cells. The expression profile of patch-seq harvested microglia in acutely isolated brain slices was correlated with the morphological and electrophysiological data. Microglia are highly dynamic immune cells that rapidly respond to an altered tissue environment resulting from traumatic injury or other pathological changes. Activation of microglial cells can already occur by the isolation procedure such as FACS sorting or tissue slicing. Morphological changes such as retraction of ramified microglial processes have been shown to occur rapidly after acute brain slicing (Matyash et al 2017). Thus, we were trying to minimize the time between tissue processing and RNA harvesting and processed samples constantly at 4°C. While it is well established that microglial cells change their membrane channel expression profile and their morphological phenotype within several hours after tissue slicing (Boucsein, Kettenmann, and Nolte 2000), we were able to analyze the cells at a time point before such changes occurred. The cells in our study had a typical membrane current profile of ramified microglia. As previously characterized, the slicing procedure leads within 12 hours to a membrane channel profile mimicking the profile of microglia as studied in tissue culture and after 24 hours as microglia in tissue culture stimulated with LPS which is due to newly expressed K^+^ channel populations (Kettenmann et al. 2011; Wolf, Boddeke, and Kettenmann 2017). The microglial cells from which we harvested the RNA did not yet show such a channel profile indicating that at least on the protein level, these activation markers were not present.

Our comparison of the expression profile of RNA isolated from microglia in the tissue context with microglia isolated as single cells from the brain tissue and subsequently purified by FACS indicated that the latter sample is characterized by a unique sub-population of microglia. This microglia cluster contains a set of signature genes associated with stress which are not found in the data obtained with the patch-seq technique. Stress adaptation responses have evolved in eukaryotic and mammalian cells in order to enable cell survival under changing conditions of the environment. In line with immediate stress responses such as post-translational modification, post-transcriptional regulation, physical regulation of ion channels and transporters, alterations to gene expression can occur within minutes after stress exposure.

In mammalian cells one of the key players among the stress-related proteins is the Activator protein-1 (AP-1) transcription factor family (Leppä et al. 1999). It comprises a number of molecules including c-Jun, c-Fos, ATF and JDP. These molecules are known to be associated with apoptosis, but also are reported to promote cell proliferation and differentiation under certain conditions. All these genes were found enriched in the FACS specific Cluster 2 and 3, but not in microglia that had been isolated by Patch-seq. Our study indicates that these genes are activated in response to the isolation and purification procedure and are not expressed in ramified (‘resting’) microglia in acute slices of the normal brain, qualifying patch-seq as a more gentle procedure for scRNA isolation than dissociation and FACS.

One of the unique and undeniable advantages of the Patch-seq method is its ability to demonstrate that transcriptional subtypes are also physiologically relevant. Transcriptomics profiles can be correlated with morphology, electrophysiological or other physiological parameters. Furthermore, the method provides data in spatial resolution. Here, however, we were not able to identify transcriptional differences between microglia in corpus callosum, hippocampus and cortex. The number of cells analyzed might be too low for a proper statement on regional differences in microglial gene expression. However, also other previous studies using alternative methods suggested that microglia are fairly homogeneous in their gene expression profiles in the normal adult mouse brain (Masuda et al. 2019). Because the Patch-seq method is rather tedious, time- and budget-consuming, compared to other single seq methods, it is not suitable for a high throughput analysis. Another major limitation of the Patch-seq method is the contamination present in the dataset, which was already previously reported by Tripathy et al for neuronal cell types (Tripathy et al. 2018). Amplification of small quantities of RNA obtained from single cells are prone to noise. Besides, the collection of cells from brain slices with a micropipette introduces additional contamination with the RNA floating in the chamber of the Patch-clamp setup and residues of damaged neuronal or glial processes which get attached to the outside of the pipette. In order to control for this contamination, we introduced multiple negative controls. As a negative control we used samples collected from the pipette with the same procedure as described for the cell collection, but omitting patching a cell. Thus we emphasize that this procedure needs to be used in combination with other scRNA-seq methods to obtain a more complete picture of the expression patter of individuall cells. Our analysis demonstrates that a defined set of genes is activated in response to the isolation and purification procedure and are not expressed in ramified (‘resting’) microglia in acute slices of the normal brain and that patch-seq is a more gentle procedure for scRNA isolation.

Studying microglial cells in their undisturbed form in the healthy brain environment is important in order to understand their physiological function. Unfortunately, most single cell -omics methods require the mechanical and/or chemical disturbance of these cells and their environment, which often times may be interpreted by microglia as a pathologic insult. It is becoming clear that tissue dissociation has a substantial impact on the transcriptome of cells as it has been revealed in different cell types (Marsh et al., 2022; van den Brink et al., 2017). This study indicates that microglia *in situ* lack transcripts associated with stress response due to tissue-dissociation. Thus, in order to get a clear picture on the transcriptomic profile of immune cells in the tissue environment, cell-activating effects due to the procedure have to be given stronger consideration during interpretation of transcriptomic data.

## STAR Methods

### ANIMALS

10-14 weeks old, male CSFR-1-EGFP (‘MacGreen’) mice (Sasmono et al. 2003; obtained from Charles River, Germany) expressing enhanced green fluorescent protein (EGFP) under control of the colony stimulating factor 1 receptor promoter were used for the study. The transgenic strain was on a C57BL/6 N genetic background. Mice were kept in the animal facility at the Max Delbrück Center under a 12-hour/12-hour dark-light cycle with food and water supply ad libitum. All experiments were performed according to the guidelines of the German law for animal protection. The animal experiments and care protocols were approved by the Landesamt für Gesundheit und Soziales, Berlin (X9005/18).

### ACUTE BRAIN SLICE PREPARATION

For preparation of acute brain slices, mice were sacrificed by cervical dislocation, mice were decapitated and the brain was immediately removed and transferred into ice-cold solution, saturated with carbogen (95% O2, 5% CO2) and containing (in mM): 230 sucrose, 26 NaHCO_3_, 2.5 KCl, 1.25 NaH2PO4, 10 MgSO_4_(7H2O), 0.5 CaCl2(2H2O) and 10 glucose. The cerebellum and the olfactory bulbs were gently removed and the brain was then fixed on the stage of a vibratome (Leica SM2000R or Leica VT1200S, Nussloch, Germany). 250μm thick coronal slices were cut between Bregma -1 to -2.5 mm to include the hippocampus and corpus callosum. Slices were kept for experiments at room temperature (RT) in gassed artificial ceribral spinal fluid (aCSF) solution for a maximum of 6h. The ACSF solution contained (mM): NaCl, 134; KCl, 2.5; MgCl_2_, 1.3; CaCl_2_, 2; K_2_HPO_4_, 1.25; NaHCO_3_, 26; D-glucose, 10; pH 7.4; with measured osmolarity of 310-320 mOsm/L.

### ELECTROPHYSIOLOGICAL RECORDINGS

Whole cell patch-clamp recordings were performed using an EPC 10 patch-clamp amplifier combined with the TIDA 5.24 software (HEKA Elektronik, Lambrecht, Germany). Acute brain slices were transferred into the holding chamber mounted on an upright microscope and were perfused with gassed ACSF at room temperature with a velocity of 3-6 mL/min. Cells were visualized with a 60x objective (Zeiss Axioskop 2 FSplus, Zeiss, Oberkochen, Germany). Micropipettes were pulled from borosilicate glass capillaries with 4-8 MΩ resistances for whole-cell patch-clamp recordings of microglial cells. The standard intracellular solution contained (mM): KCl, 130; MgCl_2_, 2; CaCl_2_ 0.5; EGTA, 5; HEPES, 10; Na-ATP, 2; pH 7.3; with measured osmolarity of 280-300 mOsm/L. 1U/μl of RNase inhibitor (RiboLock RNase-Inhibitor, Thermo Scientific) was added for better RNA recovery and 10 μM sulforhodamine 101, (SR101) (Sigma Aldrich, St. Louis, USA) for visualization of the pipette flow.

Microglial cells were identified by their EGFP fluorescence and approached by the patch pipette (**Fig.1**). Recordings with a series resistance above 65 MΩ or with unstable recording conditions were discarded. To determine the pattern of membrane currents in voltage clamp mode, a series of de- and hyperpolarizing voltage pulses were applied ranging from -160 mV to 50 mV with 10 mV increment (pulse duration 50 ms) starting from a holding of -70 mV.

### SAMPLE COLLECTION

In order to obtain high RNA recovery levels, we performed a series of initial experiments which included comparison of freshly harvested cells vs cells which were deeply frozen at -80°C in liquid nitrogen upon collection (1), different volumes of intracellular solution in the pipette (2), RLT-based modification of SMART-seq2 protocol and standard SMART-seq2 protocol (3), blocked and non-blocked TSO (4). The following protocol, which was developed after a series of optimization steps described in the Results, was used for microglia collection of all further analyzed samples. In all experimental steps, starting from transferring brain slices to the patch chamber until sequencing library preparation special care was taken to avoid any RNAse contamination: all surfaces and tubes were wiped with RNAse Away solution, gloves were worn at all times and frequently changed.

To harvest cell cytoplasm after electrophysiological recording (see above), a negative pressure of -70 mbar was applied under the control of an electronic manometer for 5-10 min. During the suction, membrane currents were closely monitored for the occurrence of leakage currents. If a leak occurred, the suction was paused and was only resumed after the patch stabilized, otherwise the cell was discarded. After the suction, the pressure was released and the patch pipette was promptly and carefully removed from the slice and from the bath. In approximately 50% of cases whole microglia somata were still attached to the pipette after withdrawal from the slice and bath. The pipette was transferred to a small tube containing 4 μl lysis buffer, the pipette tip broken at the bottom of the tube and the content of the pipette was ejected into the lysis buffer.

For negative controls the pipette was inserted into the slice the same way as described above but without patching a cell. Tubes were briefly spun down and directly underwent library preparation omitting any freezing and/or preservation.

Microglial cell collection varied between 5-10 cells per day. In total, 187 samples were collected and sequenced in 3 batches. When combined for further analysis, data was controlled for batch effect.

### SMART-SEQ2 CDNA LIBRARY PREPARATION

The samples were processed with the standard SMART-seq2 library preparation protocol, adapted from Picelli et al. (Picelli et al., 2013, 2014). In brief, after the patch clamp analysis, cells/cytosol were ejected into the tubes containing 4 μl lysis buffer, containing: Triton X-100 0.1%, dNTPs (5 mM each); Oligo-dT30VN (2.5 μM); Recombinant RNase inhibitor (1 U/μl). This mix was incubated at 72 °C for 3 min followed by 10 min at 10 °C and then was put on hold at 4 °C till the next step but no longer than for 1 h.

Next, 5.7 μl RT solution, containing Superscript II first-strand buffer (1x), 1 M Betaine, 5 mM DTT, 1 μM LNA-TSO, 6 mM MgCl2, Superscript II Reverse Transcriptase 10 U/μl, Recombinant RNase inhibitor 1.75 U/μl, was added to each sample in the lysis buffer. The mix was incubated in a thermocycler for reverse transcription with the following program:

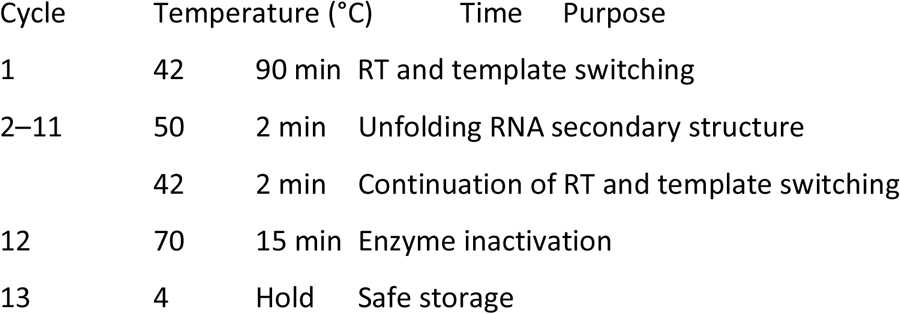

After incubation, 15 μl PCR buffer, consisting of KAPA HiFi HotStart ReadyMix (1x) and IS PCR primer 0.2 μM, was added. Next, samples were incubated with the following program:

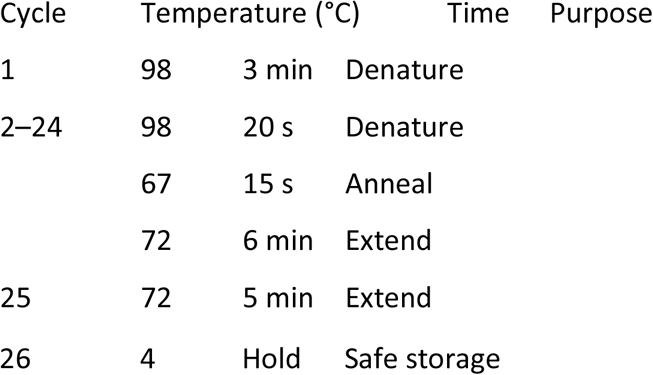

All steps of cDNA synthesis and amplification were carried out on the same day.

### NEBNEXT® ULTRATM II FS DNA LIBRARY PREPARATION AND ILLUMINA SEQUENCING

We used the NEBNext kit (NEB) for library preparation according to the manufacturer protocol, adapted to 25% reaction volume. In brief, 6μl of cDNA was fragmented by Ultra II Enzyme mix for 25 min at 37°C followed by immediate ligation with 1 μl of NEBNext Adaptor. cDNA was then cleaned up with 80% volume of AMPure XP Beads. PCR reaction with Index primers was performed of 7-9 cycles depending on initial cDNA concentration. Final libraries were purified with 80% volume of AMPure XP Beads (Beckman) and sequenced using the HiSeq4000 platform (Illumina), in paired-end sequencing mode with 2 × 75 bp read length.

### scRNA-SEQ ANALYSIS

The complete dataset consisted of three batches of samples which were collected and sequenced as described above. All the data were analyzed together. First raw reads were aligned to mouse genome using STAR aligner. We performed an assessment of the depth of sequencing and number of genes per sample. Only cell samples with more than 1000 expressed genes detected were analyzed further. There was no minimal threshold of detected genes for negative control samples. We plotted all the cell samples in the heat map for markers specific for: microglia, astrocytes, oligodendrocytes and neurons to determine the purity of the samples (Sup.Fig.1). UMAP embedding, Louvain clustering and analysis of differentially expressed genes were performed on Patch-seq samples (cells and EC) using the Seurat R toolkit version 3.6.0.

### MICROGLIA ISOLATION AND FACS

For microglia isolation we used a previously published protocol (Masuda et al., 2019). In brief, mice were perfused with PBS, brains were collected and immediately put on ice. Next, brains were cut in 8 sagittal slices on ice and placed into a glass potter. Tissue was gently homogenized with a loose pastel while being kept on ice. After washing step, suspension was mixed with 35% SIP (Stock Isotonic Percoll) and centrifuged for 25min at 4°C at 800g and stained with FACS antibodies for 30-45 min on ice. Then cell suspension was sorted on BD FACSAria™ III Cell Sorter for CD11b+/CD45low/Ly6C-/Ly6G-/PI-.

### SINGLE MOLECULE FLUORESCENT *IN SITU* HYBRIDIZATION (RNASCOPE)

*In situ* fluorescent hybridization was performed using the RNAscope Multiplex Fluorescent Reagent Kit V2 from ACDbio according to the manufacturer’s instructions. Sequences of target probes, preamplifier, amplifier, and label probe are proprietary (Advanced Cell Diagnostics, Hayward, CA). For fluorescent detection, the label probe was conjugated to Alexa Fluor 488, 546, 647, or 750 (Molecular Probes; Invitrogen, Eugene, OR).

For isolated microglial cells, cells were placed on slides using cytospin and fixed in 4% formaldehyde for 30 minutes, followed by protease digestion (2.5 μg/mL) at 23°C to 25°C. The cells were then incubated at 40°C with the following solutions: target probes in hybridization buffer A [6× SSC (1× SSC is 0.15 mol/L NaCl, 0.015 mol/L Na-citrate), 25% formamide, 0.2% lithium dodecyl sulfate, blocking reagents] for 3 hours; preamplifier (2 nmol/L) in hybridization buffer B (20% formamide, 5× SSC, 0.3% lithium dodecyl sulfate, 10% dextran sulfate, blocking reagents) for 30 minutes; amplifier (2 nmol/L) in hybridization buffer B at 40°C for 15 minutes; and label probe (2 nmol/L) in hybridization buffer C (5× SSC, 0.3% lithium dodecyl sulfate, blocking reagents) for 15 minutes. After each hybridization step, slides were washed with a wash buffer (0.1× SSC, 0.03% lithium dodecyl sulfate) two times at room temperature. For multiplex detection, equimolar amounts of target probes, preamplifier, amplifier, and label probe of each amplification system were used.

For frozen brain samples, cryosections of 12 μm thickness were dehydrated in an ethanol series, followed by protease treatment with10 μg/mL protease (Sigma-Aldrich, St. Louis, MO) at 40°C for 30 minutes in a HybEZ hybridization oven (Advanced Cell Diagnostics, Hayward, CA). Hybridization with target probes, preamplifier, amplifier, and label probe and chromogenic detection were as described above for cultured cells. We used the following probes in this study: Cdh23 (567261-C2) and Ctss (579891). Images were acquired using a Zeiss LSM 700 confocal microscope, and RNA markers were analyzed based on the average RNA dot number per cell. RNA quantity was scored based on manual counting where the positive signal is quantified as one or more dots pre cell.

### STATISTICAL ANALYSIS

Igor Pro 6.37 (WaveMetrics) and Prism 7 (GraphPad Software, San Diego, CA, USA) were used for statistical analysis in Fig. 2, 4D and Suppl. Fig. 2 and 3. Statistical significance levels are represented as *p ≤ 0.05; **P ≤ 0.01; ***P ≤ 0,001. Data are expressed as mean ± SEM.

## Acknowledgements

The authors wish to thank Caroline Braeunig, Flow Cytometry Operator for excellent technical assistance, as well as Dr. Susanne Wolf and Bilge Ugursu for helpful discussion with the FACS method; Dr. Nirmeen Elmadany for the discussions; Elijah Lowenstein for assistance with smFISH; and Dr. Christine Kocks for her valuable comments on the manuscript. This work was supported by Einstein-Stiftung, the Helmholtz-Gemeinschaft, Zukunftsthema “Immunology and Inflammation” (ZT-0027) and the Berlin Institute of Health (BIH). The authors declare no competing financial interests.

## Figure Legends

**Suppl. Fig. 1.**
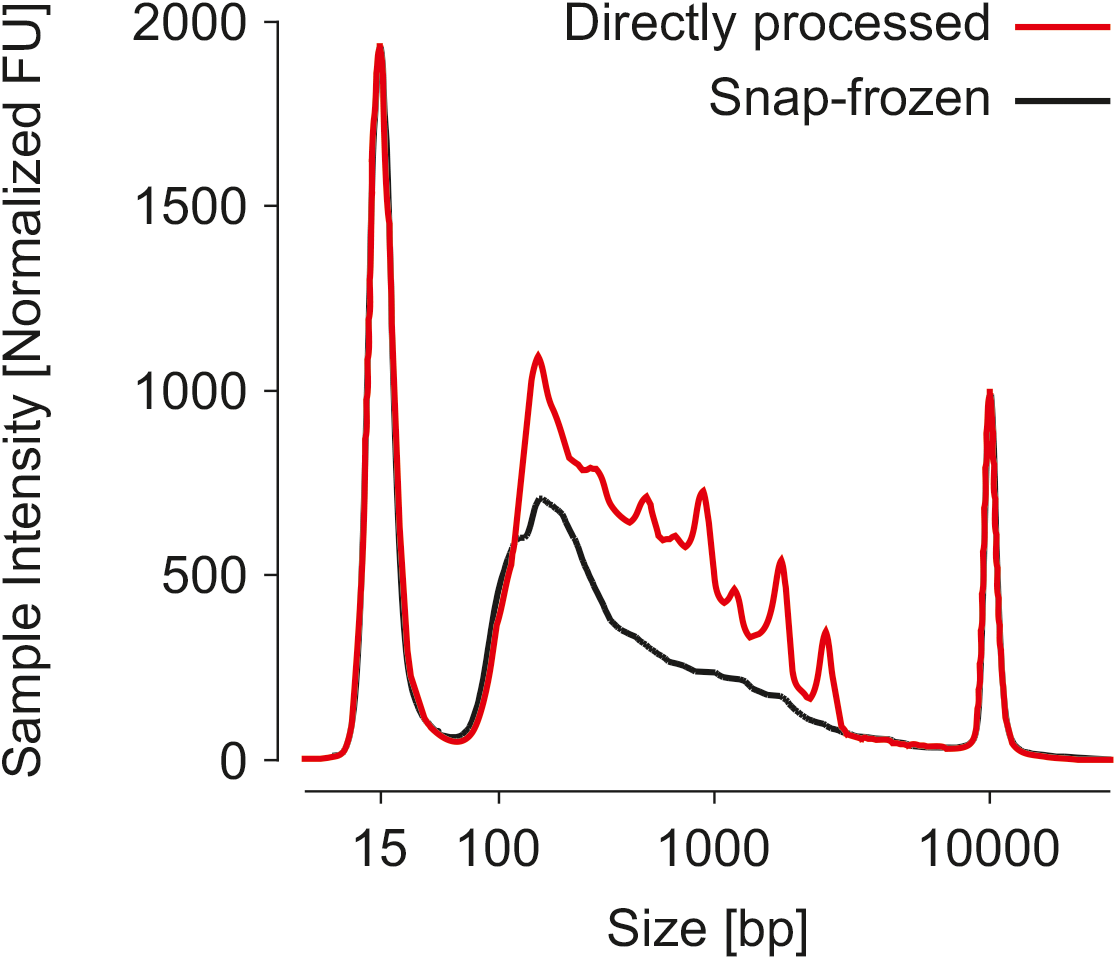
Representative tapestation profiles of microglia samples Red profile was obtained from a sample that was processed directly after harvesting the cytosol whereas the black sample was snap-frozen in liquid N_2_ for 24 hours before the SmartSeq2 protocol was applied.

**Suppl. Fig. 2.**
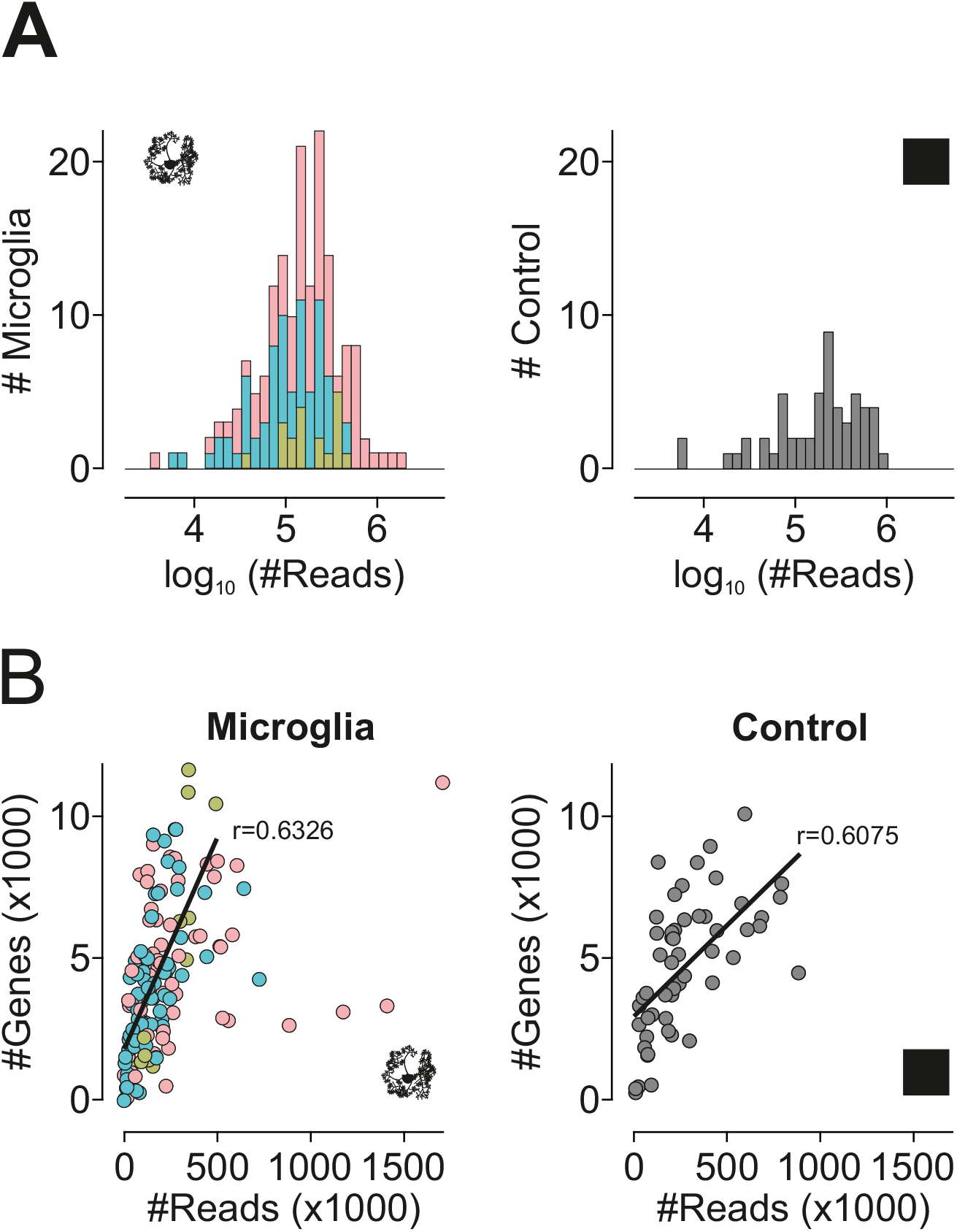
Quality control of raw sequencing data **A** Histograms depicting the number of sequenced reads obtained in MG (*left*) and EC (*right*) samples. **B** Correlation of read counts and gene numbers in MG (*left*) and EC (*right*) samples.

**Suppl. Fig. 3.**
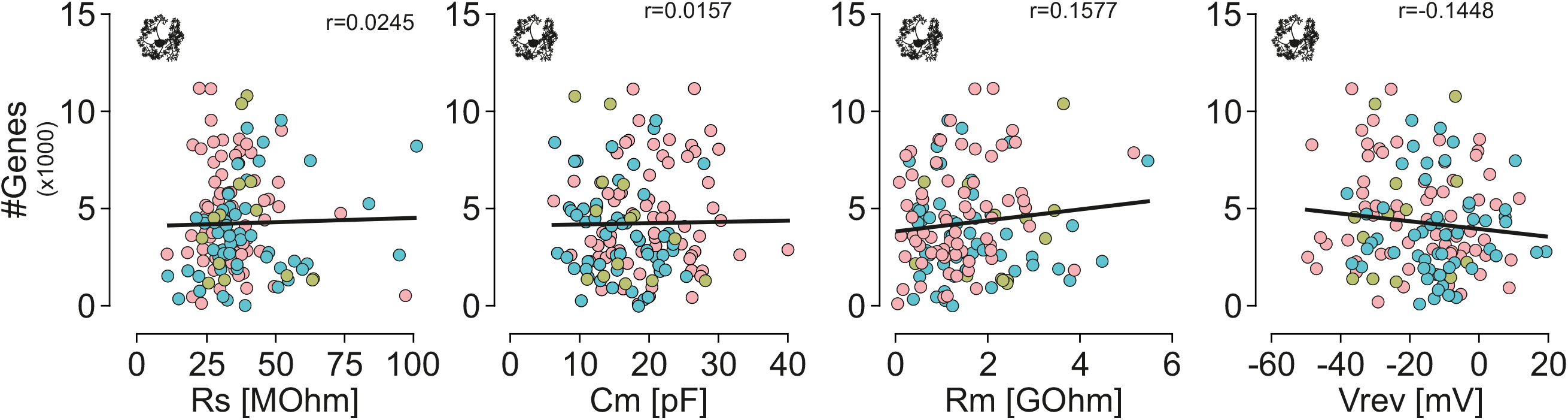
Number of genes is independent of electrophysiological properties. Correlation between series resistances (R_s_; *left*), capacities (C_m_; *middle left*), membrane resistances (R_m_; *middle right*) and reversal potentials (V_rev_; right) with gene numbers in microglia from cortex, hippocampus and corpus callosum. None of the electrophysiological properties clearly correlated with the number of genes obtained in the subsequent single cell analysis.

**Suppl. Fig. 4.**
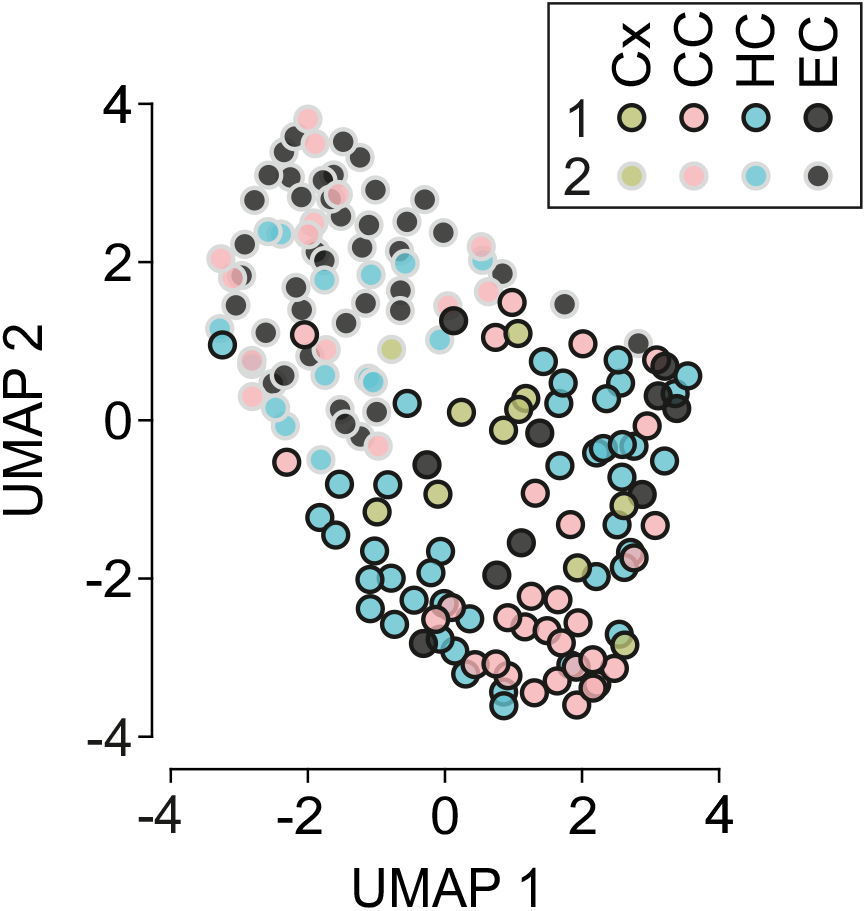
UMAP labelled by sample of origin (Variation of Fig. 3A) UMAP plot of MG (yellow=cortex; red= corpus callosum; blue= hippocampus) and EC (gray) samples. Clustering analysis revealed two different clusters which are indicated by the black (Cluster 1) and gray (Claster2) outlines.

**Suppl. Fig. 5.**
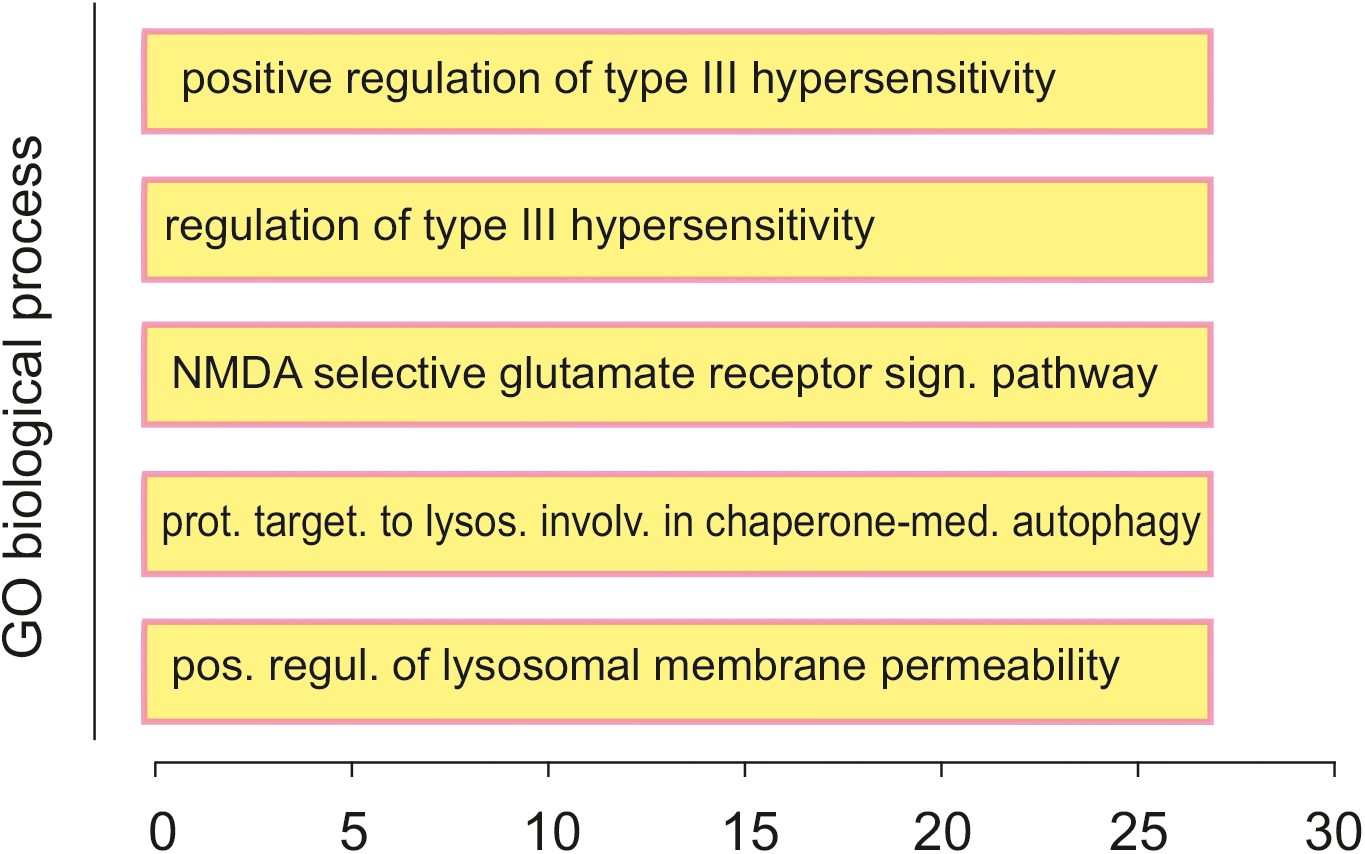
GO analysis of Clusters 1 and 4 from Fig. 4A Gene ontology (GO) enrichment analysis of marker-genes of Clusters 1 and 4 from Fig. 4A. The bar chart represents the top 5 significantly enriched pathways. X-axis shows the fold enrichment of each pathway.

